# Timing the cerebellum and its connectivity within the social brain

**DOI:** 10.1101/2024.01.09.574775

**Authors:** Andrea Ciricugno, Chiara Ferrari, Lorella Battelli, Zaira Cattaneo

## Abstract

The posterior cerebellum is a recently discovered hub of the affective and social brain, with different subsectors contributing to different social functions. However, very little is known about *when* the posterior cerebellum plays a critical role in social processing. Due to its location and anatomy, it has been difficult to use traditional approaches to directly study the chronometry of the cerebellum. To address this gap in cerebellar knowledge, here we investigated for the first time the *causal* contribution of the posterior cerebellum to social processing using a chronometric transcranial magnetic stimulation (TMS) approach. We show that the posterior cerebellum is recruited at an early stage of the emotional processing (starting from 100 ms after stimulus onset), simultaneously with the posterior superior temporal sulcus (pSTS), a key node of the emotional-social brain. Moreover, using a condition-and-perturb TMS approach, we found that the recruitment of the pSTS in emotional processing is dependent on cerebellar activation. Our results are the first to shed light on chronometric aspects of cerebellar function and its causal connectivity with other nodes of the social brain.

## Introduction

The role of the cerebellum in social cognition is well established. We can now count on solid evidence from different approaches demonstrating the role of the posterior (vs. anterior) cerebellum in a variety of social tasks, including facial and bodily emotion recognition and discrimination (Ferrari et al., 2018a, 2022a; Ferrucci et al., 2012; Fusar-Poli et al., 2009; Metoki et al., 2022), mentalizing (Clausi et al., 2022; Heleven et al., 2021; Metoki et al., 2022; Oldrati et al., 2021), biological motion perception (Ferrari et al., 2022b; Sokolov et al., 2012, 2014), and social categorization (Gamond et al., 2017). Recent evidence has also shown that the posterior cerebellum also exhibits a fine-grained functional topography (Guell et al., 2018; Guell, 2022) even within the domain of social cognition. For instance, we have recently provided causal evidence for a medial-to-lateral gradient in the posterior cerebellum, with medial cerebellar sectors supporting basic social cognitive functions (emotion recognition and processing) and more lateral/hemispheric sectors playing a role in higher-level social inferential processing and mentalizing (Ferrari et al., 2023).

Despite the extensive progress in our understanding of cerebellar topography in humans, very little is known about the *time course* of mental operations in the cerebellum. Chronometric approaches play a unique role in clarifying the cognitive operations that each cortical node within a network performs and how they are organized in real time. However, in the case of the cerebellum, chronometric approaches can be challenging to implement, mainly because the anatomy of the cerebellum makes it difficult to use traditional electrophysiological methods (i.e., electroencephalogram, EEG, and magnetoencephalography, MEG) (Andersen et al., 2020). Nevertheless, knowing *when* the posterior cerebellum is involved in different social processing stages would be crucial for a better understanding of its different functions (e.g., implementation of internal models, social sequencing, see Ito, 2008; Leggio & Molinari, 2015; Van Overwalle et al., 2019a), especially in relation to the timing of activation of other connected regions of the social brain (Sokolov et al., 2012; Van Overwalle & Mariën, 2016).

To fill this gap in current knowledge of cerebellar functions, here we systematically investigate the time course of the *causal* involvement of the cerebellum in emotional processing using a chronometric TMS approach. Indeed, thanks to its excellent temporal resolution (on the order of a few milliseconds), TMS can be used to study the chronometry of mental processes in the brain (see Cattaneo et al., 2022; de Graaf et al., 2014; Pitcher, 2014; Pitcher et al., 2007). Critically, we compared the timing of cerebellar activation with the timing of activation in other nodes of the social brain, specifically the posterior superior temporal sulcus (pSTS), and we assessed for the first time whether activation of the latter region in social processing is *causally* dependent on cerebellar activation. Although many studies have shown that the posterior cerebellum is anatomically and functionally connected to cortical hubs of the social brain, such as the pSTS and the dorsomedial prefrontal cortex (dmPFC) (Metoki et al., 2022; Sokolov et al., 2012, 2014; Van Overwalle et al., 2019b), this evidence is mostly based on fMRI studies, which cannot assess *causal* connectivity.

In the first experiment, we used TMS to investigate the chronometry of cerebellar recruitment in facial emotion discrimination, a task in which we have demonstrated a role for the posterior paravermal cerebellum (see Ferrari et al., 2018a, 2022a, 2023). We expected the cerebellum to play a role in early stages of emotional processing, in line with the idea that this region contributes to social cognition by implementing predictive models (Leggio & Molinari, 2015). In a second experiment, we causally tested the time course of pSTS involvement to compare the time course of posterior cerebellar and pSTS activation during the same emotion discrimination task. We hypothesized that these two regions work in tandem, possibly becoming active within similar time windows. Finally, in a third experiment, we used a condition-and-perturb TMS approach (a methodology based on TMS state dependence, Silvanto et al., 2008a) to directly assess causal functional connectivity between the posterior cerebellum and the pSTS. We hypothesized that if the two regions are causally functionally connected, inhibition of the cerebellum should affect responses in the pSTS.

## Experiment 1

In Experiment 1, we assessed the temporal involvement of the cerebellum in facial emotion discrimination by applying triple-pulse TMS (20 Hz) over early visual areas V1/V2 and the paravermal posterior cerebellum at four different time windows: 20-120, 120-220, 220-320, 320-420 ms from stimulus onset. V1/V2 was also targeted to rule out the possibility that cerebellar TMS effects were in fact due to the spread of activation to nearby early visual cortex (see Renzi et al., 2014). We expected cerebellar TMS to affect facial emotion discrimination in the 120-220 ms time window. Indeed, a MEG study reported cerebellar activation during passive processing of emotional scenes around 160 ms after stimulus onset (Styliadis et al., 2015), and we have previously demonstrated the causal contribution of the cerebellum in emotion discrimination tasks (similar to the one used here) using triple-pulse TMS 150 ms after the appearance of the emotional face (Ferrari et al., 2018a; 2022a, 2023).

## Method

### Participants

Twenty-five subjects (7 males, mean age = 23.9 years, SD = 2.9), with normal or corrected-to-normal vision, participated in the experiment. An a priori power analysis conducted using G-Power 3.1 software indicated that our experimental design required a sample size of at least 18 subjects to achieve 80% of power at a significance threshold alpha of 0.05, with an expected large effect size of f(U) = 0.33 (η*_p_^2^*=0.10) based on our previous data (Cattaneo et al., 2022). Prior to the experiment, participants were screened to assess their compatibility with TMS (translated from Rossi et al., 2011). The protocol was approved by the local ethics committee and participants were treated in accordance with the Declaration of Helsinki.

### Task and procedure

Stimuli consisted of images selected from the NimStim database (Tottenham et al., 2009) depicting eight male and eight female Caucasian faces (each covering approximately 23×14 degrees of visual angle) expressing the six basic emotions (happiness, sadness, surprise, fear, disgust, and anger). Stimuli were displayed on a 19’’ screen and were viewed from a distance of 57 cm. Participants were presented with pairs of emotional faces (presented sequentially and belonging to two different individuals of different gender) and had to indicate whether the two faces expressed the same emotion (two-alternative forced choice task). Each trial began with a fixation cross appearing in the center of the screen (2500 ms), followed by the presentation of the first face (500 ms), a blank screen (1000 ms), and the second face (500 ms). This was followed by a blank screen until participants responded. In half of the trials, the two faces had the same emotional expression; in the other half the two faces expressed different emotions. In each experimental block, the six emotional expressions appeared the same number of times. Participants were instructed to respond as quickly as possible by pressing the left or right arrow key with their right hand (response key assignment was counterbalanced across participants). Participants performed the emotion discrimination task three times, one for each TMS site (the cerebellum, V1/V2, and the vertex), in a counterbalanced order. Each block consisted of 72 trials (18 for each TMS interval). The experimental blocks were preceded by a short practice block (10 trials) without TMS. The software E-prime 2.0 (Psychology Software Tools, Pittsburgh, PA) was used for stimulus presentation, data collection, and TMS delivery. Each session lasted on average 1 h and 30 min (including instructions, completion of the TMS questionnaire and informed consent, neuronavigation to identify the cortical hotspot, and debriefing).

### TMS

TMS was applied using a Magstim Rapid^2^ stimulator (Magstim Co, Ltd, Whitland, UK) connected to a 70-mm butterfly coil. Prior to the experiment, single pulse TMS was applied over the left motor cortex (M1) at increasing intensities to determine each participant’s resting motor threshold (rMT). Individual rMT was defined as the minimum intensity of the stimulator output that produced motor-evoked potentials (the motor response measured by electrodes placed on the hand muscles) with an amplitude of at least 50 mV in the first dorsal interosseous with a 50% probability (Rossini et al., 1994; see also Hanajima et al., 2007 for methodological details of this standard procedure). Participants were stimulated at 100% of their rMT, as in previous TMS studies targeting the cerebellum (e.g., Demirtas-Tatlidede et al., 2011; Ferrari et al., 2018a, b; 2022a, b, 2023). The average stimulation intensity was 49.80% of the maximum stimulator output (SD=2.58) and it was kept constant for the stimulation of all the target sites. Triple-pulse 20 Hz TMS was delivered over the paravermal cerebellum, V1/V2, and the vertex (control site) at four different latencies after the onset of the second face: 20-120, 120-220, 220-320, or 320-420 ms.

To localize the cerebellar targets for each individual we used a stereotaxic navigation system and the magnetic resonance images (MRIs) obtained by a 3D warping procedure that fitted a high-resolution MRI template with the participant’s scalp model and craniometric points (Softaxic 3.0, EMS, obtained using individual estimated MRI scans, see Carducci and Brusco, 2012). Anatomical Talairach coordinates (Talairach and Tournoux, 1988) of the targeted paravermal cerebellum (x=-15, y=-82, z=-32) were taken from a previous neuroimaging study of facial emotional processing (Schraa-Tam et al., 2012) and correspond to the left medial sector of lobule VI/VII. In previous studies, we demonstrated that targeting this region with TMS significantly impairs facial emotional processing (Ferrari et al., 2018a; 2021, 2023). The vertex was localized as the point halfway between the nasion and the inion on the same midline. Early visual cortex (V1/V2) was functionally localized in each participant using visual phosphenes (see Walsh and Pascual-Leone, 2003). Specifically, the coil is initially positioned 2 cm above the inion and its position is subsequently adjusted until foveal phosphenes are induced. Phosphenes were searched for each participant after dark adaptation using a modified binary search algorithm that defines the phosphenes threshold (PT) (Tyrrell and Owens, 1988; Thilo et al., 2004). The mean PT was 65.26% (SD = 13.65). For participants who could not perceive phosphenes (N = 6), the early visual cortex was localized as the point lying 1.5 cm superior to the inion on the midline connecting the inion to the nasion (Campana et al., 2006; Cattaneo et al., 2014; Heinen et al., 2005) and further confirmed by neuronavigation. For cerebellar and V1/V2 stimulation, the coil was placed tangential to the scalp and held parallel to the midsagittal line with the handle pointing superiorly, consistent with evidence that this is an effective coil orientation for modulating activity in cerebellar and visual structures (e.g. Bijsterbosch et al., 2012; Kanai et al., 2011; van Dun et al., 2017). For the vertex stimulation, the coil was placed tangentially to the scalp and held parallel to the midsagittal line with the handle pointing backwards. No participant reported any discomfort or adverse effects during TMS.

## Results

Deidentified data for all experiments are available at Zenodo.org. Mean accuracy rates and mean correct reaction times (RTs, recorded from the *onset* of the second face) were computed for each participant in each experimental condition. Accuracy scores and RTs were analyzed using a repeated measures ANOVA, with TMS site (vertex, V1/V2, cerebellum) and TMS time (20-120, 120-220, 220-320, 320-420 ms from the onset of the second face) as within-subjects variables.

Analysis of mean accuracy rates revealed no significant main effect of either TMS site, *F*(2,48)<1, *p*=.44, or TMS time, *F*(3,72)<1, *p*=.53. The interaction TMS site x TMS time, *F*(6,144)=2.99, *p*=.009, η*_p_^2^* =.11, reached significance and was further examined by looking at the simple main effect of TMS site within each time window. The analysis revealed a significant main effect of TMS site for the 20-120 ms window, *F*(2,48)=3.56, *p*=.036, η*_p_^2^* =.13, and for the 120-220 ms window, *F*(2,48)=4.55, *p*=.016, η*_p_^2^* =.16, whereas no significant effect of TMS was observed for either the 220-320 ms window, *F*(2,48)=1.13, *p*=.33, or in the 320-420 ms window, *F*(2,48)<1, *p*=.40. Post-hoc *t-*tests showed that within the 20-120 ms time window, TMS over V1/V2 impaired performance relative to the vertex, *t*(24)=2.51, *p*=.019 (*p*=.057 Bonferroni-Holm corrected), and relative to the cerebellum (although in this case the difference only approached significance), *t*(24)=2.01, *p*=.056 (*p*=.11 Bonferroni-Holm corrected). In contrast, TMS over the cerebellum did not affect accuracy compared to the control vertex condition, *t*(24)<1, *p*=.93. For the 120-220 ms time window, accuracy rates were lower when TMS was applied over the cerebellum compared to the vertex, *t*(24)=3.11, *p*=.005 (*p*=.014 Bonferroni-Holm corrected) and V1/V2, *t*(24)=2.51, *p*=.019 (*p*=.038 Bonferroni-Holm corrected) (Figure 2). No difference was observed between TMS over the vertex and V1/V2, *t*(24)<1, *p*=.76.

**Figure 1.**
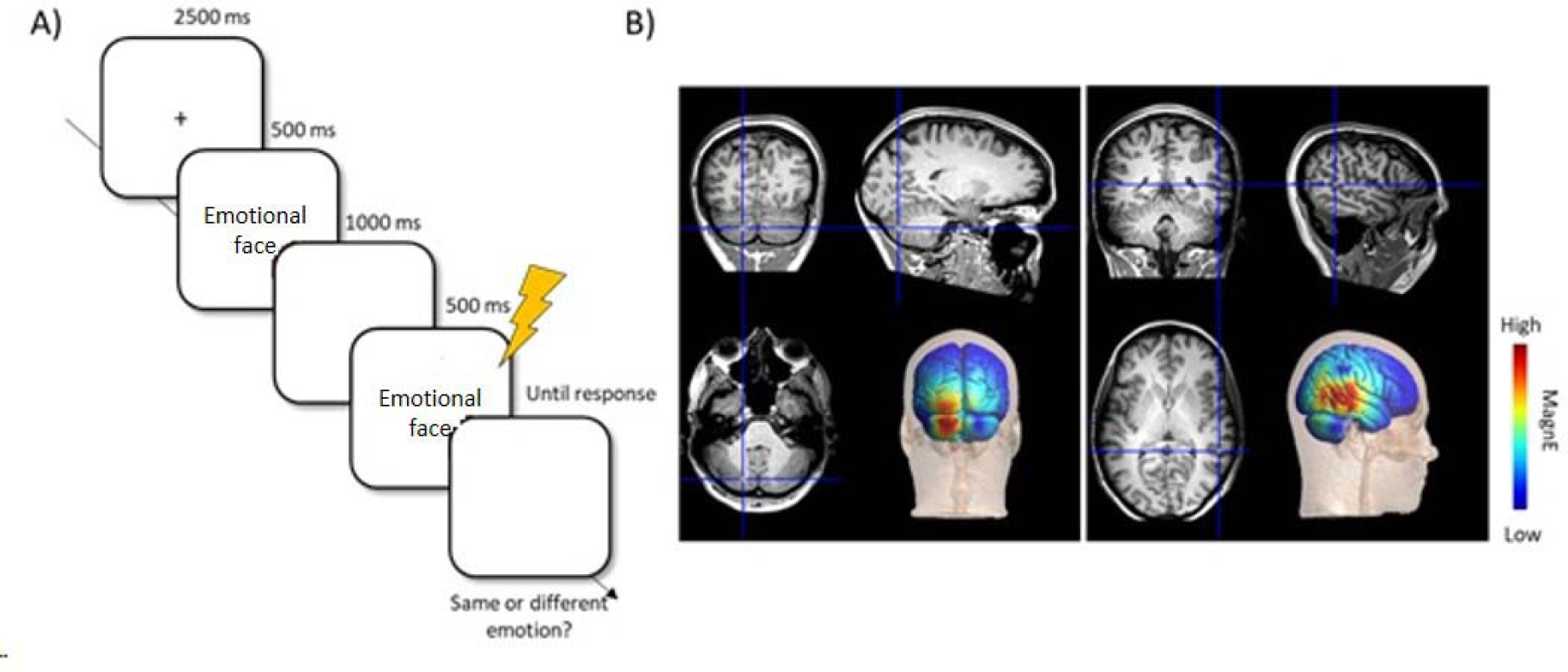
**A)** Timeline of an experimental trial of the emotion discrimination task. Each trial started with a fixation cross (2500 ms), followed by the first face (500 ms), a blank screen (1000 ms), and the second face (500 ms). Participants had to indicate whether the two faces expressed the same or a different emotion (by left/right key pressing). **B)** Coronal, sagittal and axial views (from upper left in clockwise order) of a representative participant showing the localization of the regions targeted in the present study: posterior cerebellum (left panel: TAL; x=-15, y=-82, z=-32) and pSTS (right panel: TAL: x=52, y=-48, z=8). In the lower right of each panel a representation of the estimated electric field induced by the Magstim Rapid2 stimulator 70-mm figure-of-eight coil obtained using SimNIBS (Thielscher et al., 2015; Weise et al., 2020).

**Figure 2.**
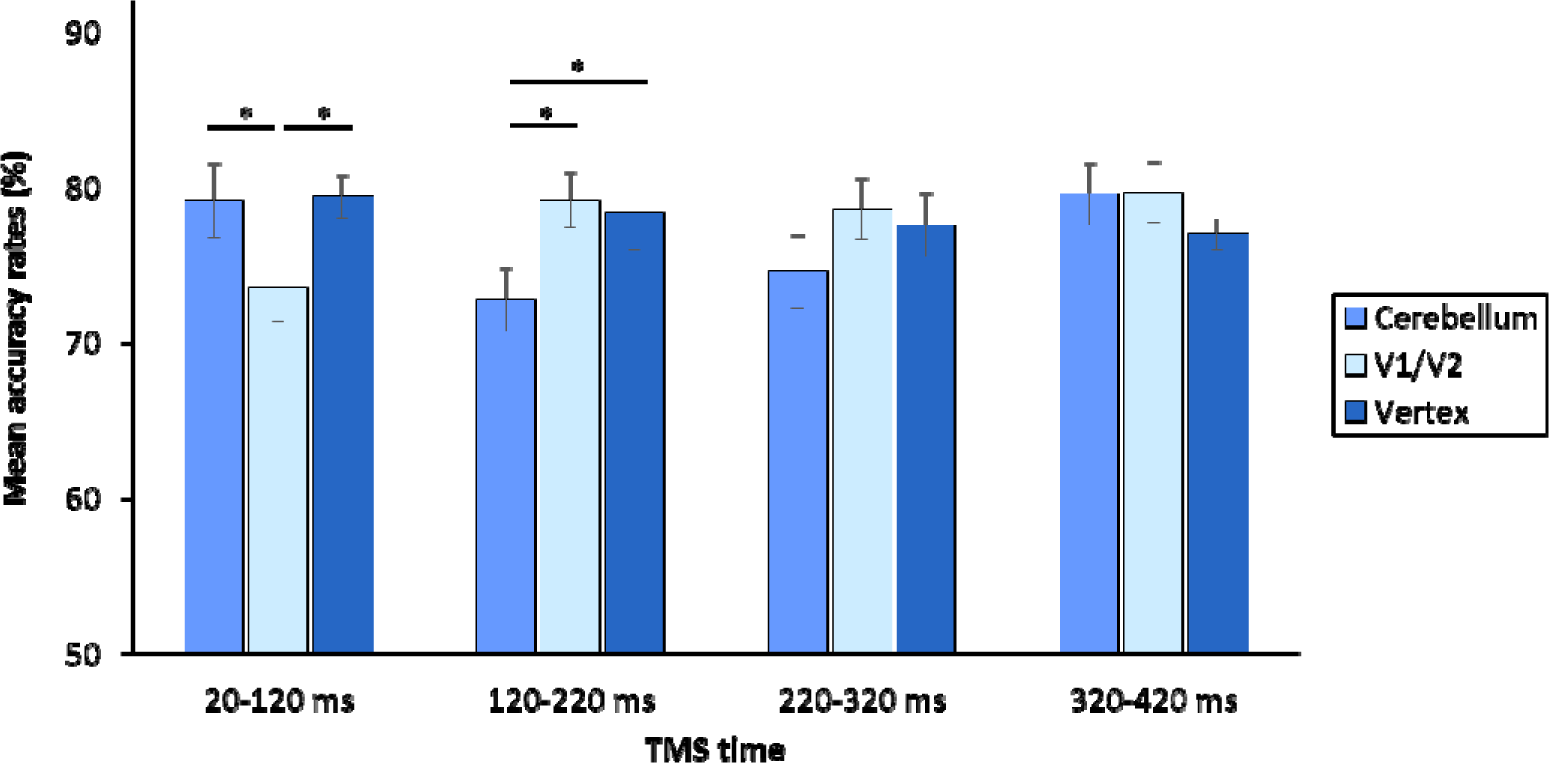
Mean accuracy rates (%) as a function of TMS site (Vertex, V1/V2, cerebellum) and TMS time (20-120 ms, 120-220 ms, 220-320 ms, 320-420 ms). Error bars indicate ±1 SEM. Asterisks indicate significant differences (p<.05, Bonferroni-Holm corrected; note the difference between TMS over V1/V2 and the cerebellum within the 20-120 ms time window only approached significance, *p*=.056 uncorrected) across conditions.

Mean correct RTs were 1121 ms (SD=249) for vertex TMS, 1135 ms (SD=344) for cerebellar TMS, and1048 ms (SD=270) for V1/V2. Analyses on RTs revealed no significant main effects or interactions: TMS site, *F*(2,48)=1.27, *p*=.29; TMS time, *F*(3,72)<1, *p*=.63; TMS site by TMS time, *F*(6,144)<1, *p*=.55.

## Experiment 2

In Experiment 1, we showed that TMS over V1/V2 disrupted the early stages of perceptual processing (20-120 ms time window), whereas cerebellar TMS affected performance at a later stage (120-220 ms), suggesting that cerebellar TMS effects are unlikely to be due to spread of activation from the nearby visual cortex (de Graaf et al., 2014). In Experiment 2, we performed a finer temporal analysis of the cerebellar contribution to facial emotion processing, focusing on the 120-220 ms time window identified in Experiment 1. Specifically, we delivered double-pulse TMS at 100-140 ms, 130-170 ms, and 170-210 ms from face onset. A previous TMS study using a similar protocol (i.e., dual-pulse TMS, same time windows, Pitcher, 2014) found that pSTS was causally involved in facial emotion discrimination in the 100-140 ms time window. Furthermore, a dynamic causal modeling (DCM) study has suggested that during social tasks (i.e., biological motion discrimination), visual input first reaches pSTS and then the cerebellum, which then sends feedback projections to pSTS, thus the two regions appear to be bidirectionally connected (Sokolov et al 2012).

## Method

### Participants

Twenty-four subjects (9 males, mean age = 22.30 years, SD = 2.35), with normal or corrected-to-normal vision participated in Experiment 2 (sample size estimation was calculated as in Experiment 1). None of them participated in Experiment 1. Prior to the experiment, participants were screened to assess their compatibility with TMS (translated from Rossi et al., 2011). The protocol was approved by the local ethics committee and participants were treated in accordance with the Declaration of Helsinki.

### Task procedure and TMS

Task and procedure were identical to Experiment 1. Regarding the TMS procedure, in Experiment 2, we delivered double-pulse TMS over the cerebellum, the vertex, and pSTS at 3 different latencies: 100-140, 130-170, and 170-210 ms from the onset of the second face of the emotion discrimination task. Double-pulse TMS and time windows were chosen based on a previous chronometric study by Pitcher (2014), which assessed the time course of pSTS involvement in facial emotion discrimination. The right pSTS target site was localized by neuronavigation (see Experiment 1); the Talairach coordinates (x=52, y=-48, z=8) were taken from a previous fMRI study that reported stronger activation of the right pSTS when individuals viewed emotional faces compared to neutral faces (Engell & Haxby, 2007; see also Ferrari et al., 2018b for 9 a TMS study using the same coordinates). For the pSTS stimulation, the coil was held parallel to the midsagittal line. The cerebellar target site and the vertex were localized as in Experiment 1. The stimulation intensity was set at 100% of rMT (mean rMT was 49.30%, SD = 2.95).

## Results

Accuracy scores and RTs for correct responses were analyzed using separate repeated measures ANOVAs, with TMS site (vertex, cerebellum, pSTS) and TMS time (100-140, 130-170, 170-210 ms) as within-subject variables.

Analyses of mean accuracy rates revealed a significant main effect of TMS site, *F*(2,46)=4.93, *p*=.012, η*_p_^2^* =.18, indicating that TMS over both the pSTS, *t*(23)=3.13, *p=*.005 (*p*=.014 Bonferroni-Holm corrected), and the cerebellum, *t*(23)=2.20, *p*=.038 (*p*=.077 Bonferroni-Holm corrected) impaired performance compared to TMS over the control site (vertex). TMS over the pSTS and the cerebellum similarly impaired performance, *t*(23)=1.03, *p*=.32 (Figure 3). The main effect of TMS time, *F*(2,46)<1, *p*=.92, and the interaction of TMS site by TMS time, *F*(4,92)<1, *p*=.82, were not significant.

**Figure 3.**
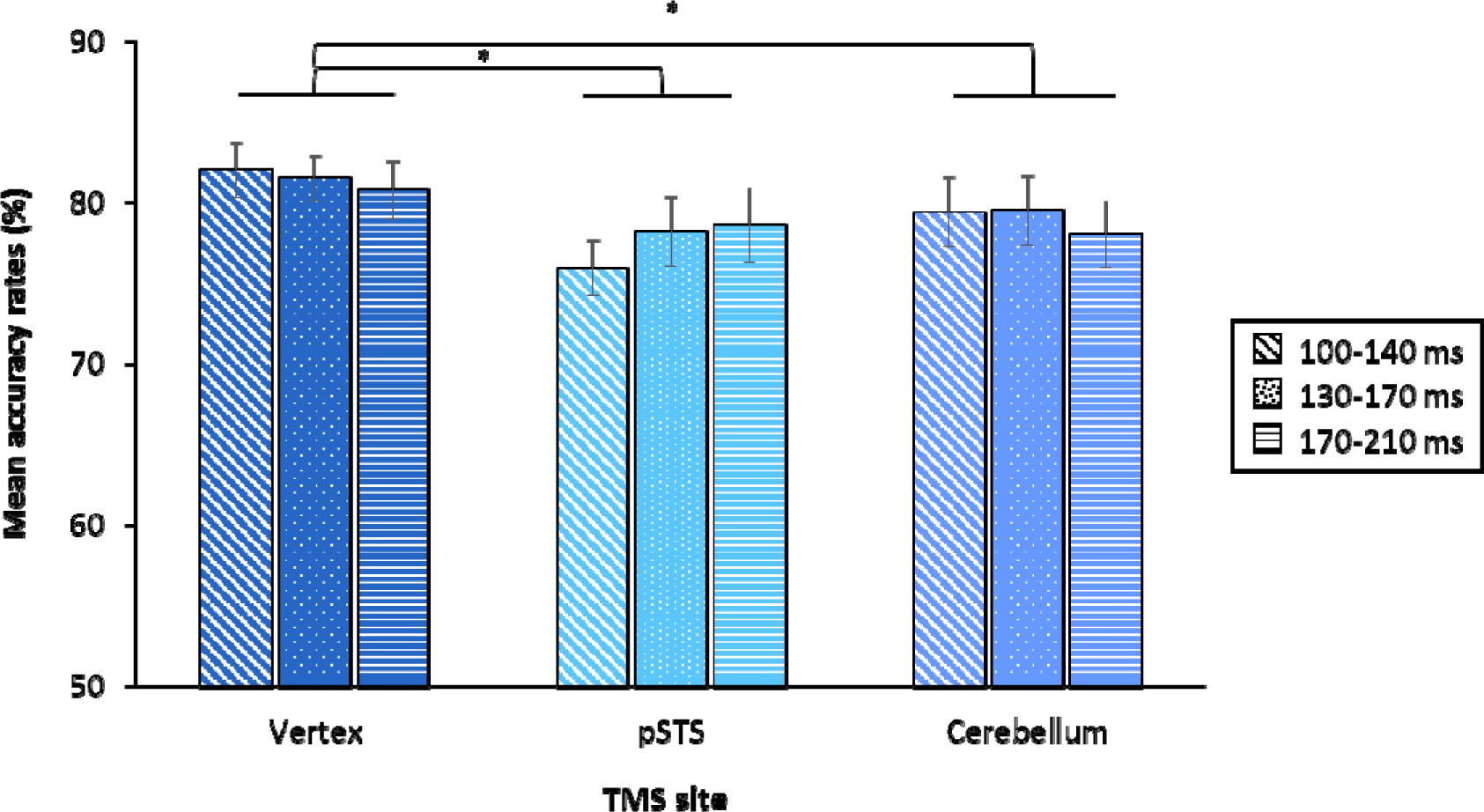
Mean accuracy rates (%) as a function of TMS site (Vertex, pSTS, Cerebellum) and TMS time (100-140 ms, 130-170 ms, 170-210 ms). Error bars indicate ±1 SEM. Asterisks indicate significant differences (p<.05, Bonferroni-Holm corrected) across conditions.

Mean correct RTs were 1074 ms (SD=200) for vertex TMS, 1084 ms (SD=256) for cerebellar TMS, and 1076 ms (SD=280) for pSTS TMS. The ANOVA on RTs revealed no significant main effects or interactions: TMS site, *F*(2,46)<1, *p*=.98; TMS time, *F*(2,46)<1, *p*=.53; TMS site by TMS time, *F*(4,92)=1.46, *p*=.22.

## Experiment 3

In Experiment 2, we showed that TMS over the cerebellum and over the pSTS affected performance similarly across all the time windows we tested. Thus, our data suggest that both the pSTS and the posterior cerebellum are *causally* involved in emotion discrimination from 100 ms after stimulus onset. We know from neuroimaging evidence that these regions co-activate in several social tasks (e.g., Sokolov et al., 2012), and a common hypothesis is that these two regions communicate bidirectionally (Sokolov et al., 2012). If so, perturbing activity in one of these regions should affect information processing in the other (Silvanto et al., 2008a, b). To directly test this hypothesis, in Experiment 3 we used a dual-site condition-and-perturb TMS approach to investigate whether the application of inhibitory 1Hz TMS over the cerebellum modulates pSTS recruitment during emotion discrimination. If these regions are functionally connected, this should be the case.

## Method

### Participants

Thirty-eight healthy volunteers (10 males, mean age = 23.19 years, SD = 3.08), with normal or corrected-to-normal vision, were recruited to participate in the experiment. None of them participated in the two previous experiments. An a priori power analysis performed using G-Power 3.1 software indicated that our experimental design required a sample size of 38 individuals to achieve 80% of power at a significance threshold alpha of 0.05, with an expected large effect size of f(U) = 0.48 (η*_p_^2^* = 0.19) based on previous data (Slivinska et al., 2020). Prior to the experiment, participants were screened to assess their compatibility with TMS (translated from Rossi et al., 2011). The protocol was approved by the local ethics committee and participants were treated in accordance with the Declaration of Helsinki.

### Procedure

Participants completed two experimental sessions on separate days (session duration was kept constant for each individual; mean interval between sessions = 8 days; minimum interval = 5 days). At the beginning of each session, single pulse TMS was applied to determine each participant’s rMT (measured as in the previous experiments). Mean rMT was 49.6% (SD = 2.9) and 49.5% (SD = 2.6) for the first and second sessions, respectively. A paired-sample t-test revealed no difference in rMT between the two sessions (t(35)<1, p=.52). Participants were administered 15 minutes of conditioning (offline) 1 Hz TMS over the posterior cerebellum or the vertex (control conditioning condition) and then performed two blocks of the emotion discrimination task used in the previous experiments while receiving online TMS over the right pSTS or the vertex (control perturbation condition) (see Figure 4). Specifically, the conditioning rTMS consisted of 900 pulses at 1 Hz delivered offline (i.e., before the task) at 100% of participants’ rMT, consistent with previous TMS studies targeting the cerebellum (e.g. Demirtas-Tatlidede et al., 2011; Ferrari et al., 2018a, 2021, 2023). 1 Hz rTMS is thought to reduce the neural excitability of the targeted region and to subsequently disrupt its associated functions for at least half of the stimulation duration (Walsh & Pascual-Leone, 2003). Following conditioning TMS, participants performed two blocks of the emotion discrimination task (each composed of 40 trials), one for each TMS site (pSTS and vertex). During the task, at the onset of the second face (target image), participants received trains of five 10 Hz TMS pulses (i.e., one pulse every 100 ms) at 120% of rMT. These stimulation parameters were chosen to maximize the disruptive effect of TMS, consistent with previous studies targeting pSTS (Sliwinska & Pitcher, 2018; Sliwinska et al., 2020). Each block lasted approximately 3.5 min ensuring that the task was performed within a time window in which offline TMS is effective (Battelli et al., 2009, 2017). The order of stimulation sites for both conditioning and perturbing TMS was counterbalanced across subjects. The target sites were localized as in the previous experiments. Each session lasted an average of 1 hour and 30 min (including instructions, completion of the TMS questionnaire and informed consent, neuronavigation, and debriefing). No participant reported any discomfort or adverse effects during TMS.

**Figure 4.**
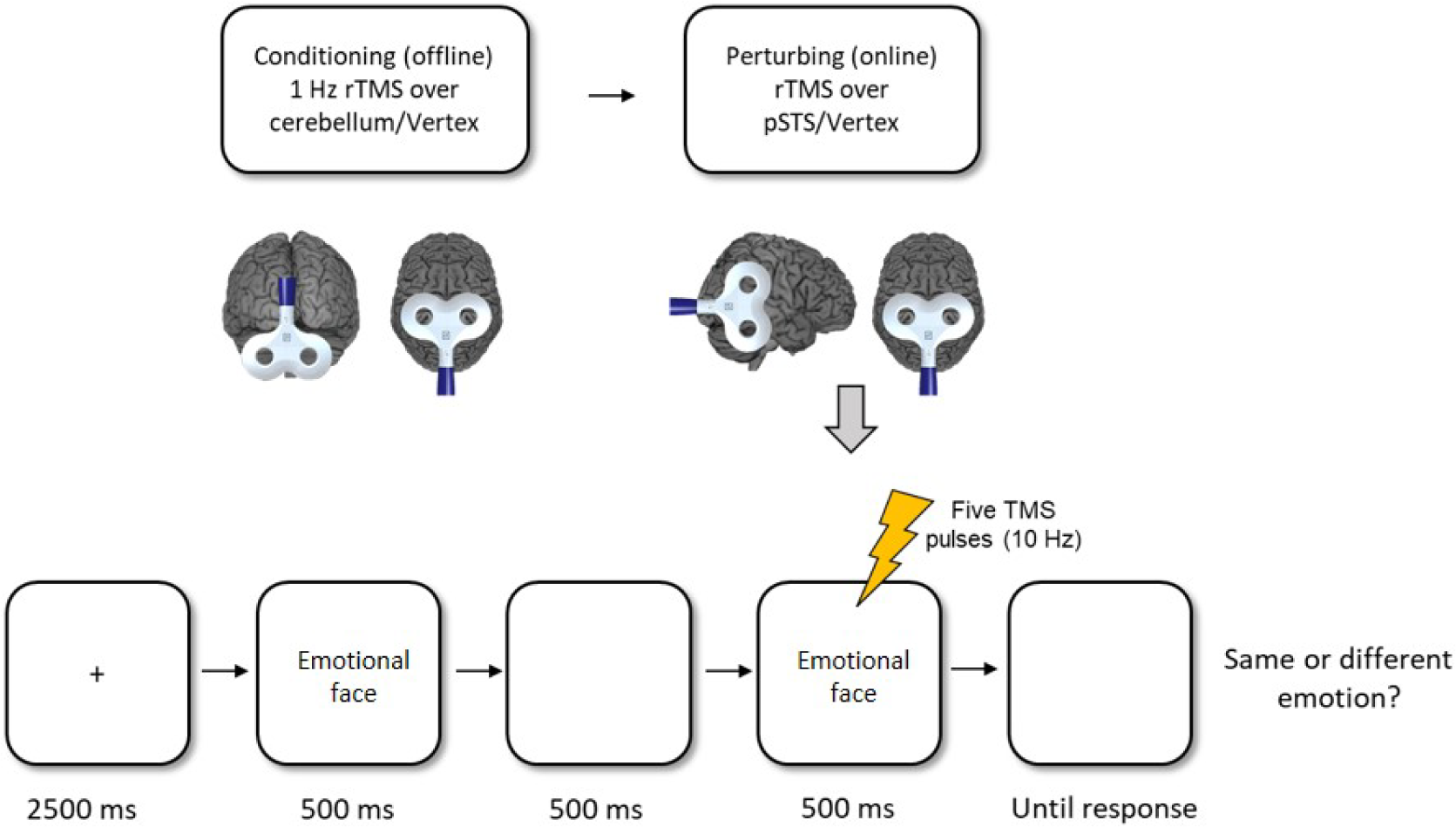
Timeline of an experimental session. Participants received 15 minutes of 1 Hz (offline) conditioning rTMS over the paravermal cerebellum and the vertex (in a counterbalanced order in two experimental sessions performed on separate days). Following conditioning rTMS, participants performed the facial emotion discrimination task (the same used in Experiments 1 and 2) while receiving perturbing (online) rTMS over the pSTS and the vertex (in a counterbalanced order).

## Results

Deidentified data for all experiments are available on Zenodo.org. Data from two participants were excluded due to technical problems. Mean accuracy rates and mean correct RTs (recorded from the onset of the second face of the emotion discrimination task) were computed for each participant in each experimental condition. Accuracy scores and RTs were analyzed using separate repeated measures ANOVAs, with conditioning TMS (cerebellum and vertex) and perturbing TMS (pSTS and vertex) as within-subject variables.

Analysis of mean accuracy rates revealed no significant main effect for conditioning TMS, *F*(1,35)<1, *p*=.65. The main effect of perturbing TMS was significant, *F*(1,35)= 9.89, *p*=.003, η*_p_^2^* =.22, as was the interaction of perturbing TMS by conditioning TMS, *F*(1,35)=13.22, *p<*.001, η*_p_^2^* =.27 (Figure 5).

**Figure 5.**
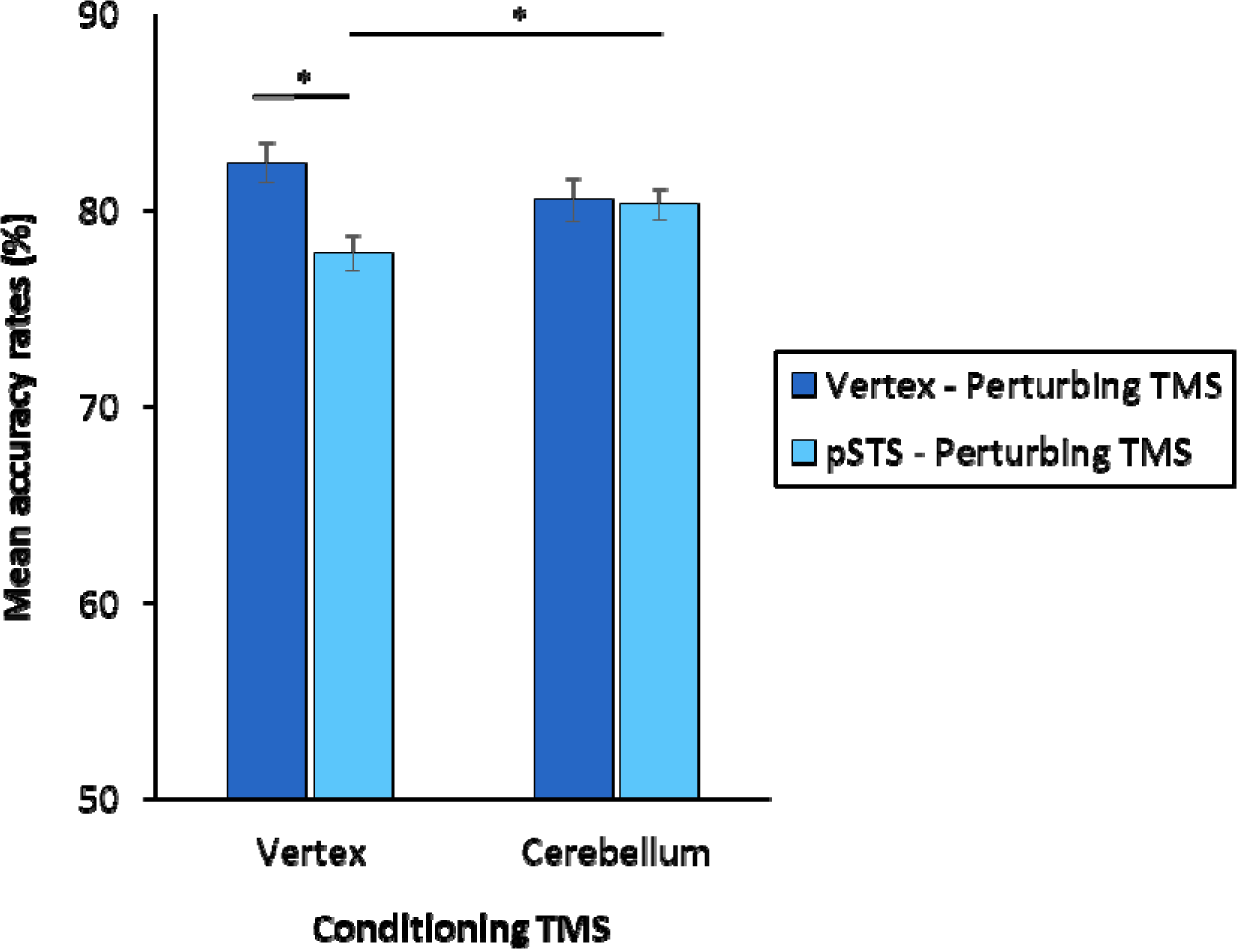
Mean accuracy rates (%) as a function of conditioning TMS (vertex, cerebellum) and perturbing TMS (vertex, pSTS). Error bars indicate ±1 SEM. Asterisks indicate significant differences (p<.05, Bonferroni-Holm corrected) across conditions.

Post-hoc comparisons showed that when online TMS was applied over the vertex, performance after 1 Hz conditioning of the cerebellum tended to be lower compared to conditioning of the vertex, *t*(35)=1.88, *p*=.068 (*p*=.14 Bonferroni-Holm corrected). Furthermore, following 1 Hz vertex conditioning, participants’ accuracy was lower when we delivered online TMS over the pSTS (mean=78%, SD=6.6) compared to the vertex (mean=82%, SD=6.1), *t*(35)=4.07, *p<.*001, *d*=.68 (*p*=.002 Bonferroni-Holm corrected), indicating effective modulation of pSTS activity. Critically, this pattern disappeared following 1 Hz conditioning of the posterior cerebellum: in this condition perturbing pSTS activity with online TMS (mean=80%, SD=4.8) did not affect performance compared to online TMS of the vertex (mean=81%, SD=5.4), *t*(35)<1, *p*=.82. Accordingly, performance was higher when pSTS was stimulated online following cerebellar conditioning compared to vertex conditioning, *t*(35)=2.58, *p*=.014 (*p*=.042 Bonferroni-Holm corrected).

Mean correct RTs after vertex conditioning TMS were=996 ms (SD=326) for perturbing vertex TMS and 976 ms (SD=332) for perturbing pSTS TMS. Mean correct RTs after cerebellar inhibitory conditioning TMS were 959 ms (SD=210) for perturbing vertex TMS and 960 ms (SD=218) for perturbing pSTS TMS. Analysis on RTs revealed no significant main effects or interactions: perturbing TMS, *F*(1,35)<1, *p*=.64; conditioning TMS, *F*(1,35)<1, *p*=.50; conditioning TMS by perturbing TMS, *F*(1,35)<1, *p*=.45.

## Discussion

In this study, we conducted the first investigation of the time course of the *causal* recruitment of the posterior cerebellum in social processing, taking into account the time course of recruitment of other nodes of the social brain (i.e. the pSTS), and we assessed causal functional connectivity between the cerebellum and the pSTS by using a perturb-and-measure TMS approach.

In Experiment 1, we applied repetitive TMS in four different 100ms-long time windows (starting at 20, 120, 220, and 320 ms from stimulus onset). Our results showed that cerebellar TMS significantly impaired emotion discrimination when administered in the 120-220 ms time window, but not at other time points. This result provides the first *causal* evidence of the time course of the posterior cerebellum’s involvement in a social task and, more broadly, in cognitive processes. Importantly, these effects were not due to indirect stimulation of V1/V2, as direct stimulation of these areas (as expected, see de Graaf et al., 2014) only affected performance in the earliest time window (20-120 ms).

In Experiment 2, we conducted a more detailed temporal analysis of the role of the cerebellum in facial emotion processing using double-pulse TMS, and focusing on shorter 40 ms time windows (as in Pitcher, 2014). We also compared the effects of cerebellar stimulation with those of stimulation of the pSTS, a critical region of the social brain critical for facial emotion processing. Our results demonstrated that TMS applied to both regions significantly affected performance at all time windows we investigated (100-140 ms, 130-170 ms, 170-210 ms). Previous MEG studies reported a strong pSTS response up to 250 ms after stimulus onset during emotion judgment tasks involving facial expressions (Seitz et al., 2008; Streit et al., 2003), whilst previous TMS evidence observed a specific contribution of the pSTS to emotion discrimination within the time window of 60 to 140 ms after stimulus onset (Pitcher, 2014).

Recruitment of the pSTS in facial emotion discrimination has been interpreted as the rapid integration of information derived from changeable aspects of the stimuli (e.g., facial expressions, movements) to attribute social and emotional meaning to faces and thereby understand the intentions of others (Walbrin et al., 2018; Zhu et al., 2013). These operations represent an intermediate step between basic perceptual mechanisms and higher-level social processes (Santavirta et al., 2023; Zhu et al., 2013). While the data from Experiment 2 are consistent with previous evidence regarding the pSTS, the critical finding here is that the posterior cerebellum causally contributes to emotion discrimination within the same time window as the pSTS. This suggests a collaborative effort between these two regions in supporting this cognitive process.

A dynamic causal modeling (DCM) study has proposed that during social tasks, such as in biological motion processing, the visual input travels from extrastriate visual regions to the pSTS (see e.g., Babo-Rebelo et al., 2022; Ethofer et al. 2011; Pitcher and Ungerleider, 2021; see also Borgomaneri et al., 2023 for evidence on the role in social processes of reentrant pSTS-to-visual cortex back-projections). From there, it travels from the pSTS, via the pons, to the lateral cerebellum (which does not receive direct input from the visual cortices, see Buckner, 2011). The cerebellum then sends feedback projections, via the thalamus, to the pSTS, making the two regions appear to be bidirectionally connected (Sokolov et al 2012). Within this circuit, the role of the cerebellum may be to fine-tune the contribution of the pSTS to mental state inference by constructing situational internal models. Indeed, a consistent body of evidence suggests that the cerebellum anticipates the behavior of others by generating internal models of social interactions that predict potential outcomes based on individuals’ prior experience and knowledge (Ito, 2008; Leggio and Molinari, 2015; Sokolov et al., 2017; Van Overwalle et al., 2020). In Experiment 2, we demonstrated for the first time that, if this is indeed the case, the paravermal cerebellum generates these internal models for emotion discrimination around 100 ms from stimulus onset, concurrently with the operations performed by the pSTS.

It could be argued that the simultaneous causal recruitment of the posterior cerebellum and the pSTS may reflect the simultaneous activation of two distinct networks rather than an effective functional interaction between these two regions, as part of the same network. In Experiment 3, we ruled out this possibility. Specifically, we showed that the recruitment of the pSTS in face emotion recognition is dependent on cerebellar activation. Indeed, disrupting pSTS activity with online (i.e., during the task) repetitive TMS impaired participants’ performance in discriminating emotional face expressions, in line with previous studies (Pitcher, 2014; Pitcher et al., 2007; Sliwinska et al., 2020; Sliwinska and Pitcher 2018). Crucially, however, when we inhibited posterior cerebellar activity with offline 1 Hz TMS prior to the task, online perturbation of pSTS activity no longer impaired performance. This result demonstrates that the recruitment of the pSTS during facial expression discrimination depends on the level of cerebellar activation. Thus, our data demonstrate for the first time, the existence of *causal* functional connections between the posterior cerebellum and pSTS during social processes. This is consistent with neuroimaging DCM data supporting the existence of effective bidirectional connectivity between the left cerebellum and the right pSTS during social tasks, such as biological motion perception (Sokolov et al., 2012). In addition, anatomical studies have shown that the cerebellum is structurally connected to the pSTS (Schmahmann & Pandya, 1991; Sokolov et al., 2014).

Data from Experiment 3 are likely indicative of state-dependent effects of TMS (Silvanto et al., 2017, 2018). State dependence in TMS studies refers to the consistent observation that when the activity of a brain region is primed by inhibitory offline 1Hz rTMS, subsequent online TMS applied over the same region either has no effect or can even paradoxically cause a behavioral facilitation, as opposed to the disruptive effect typically observed when online TMS alone is administered. In other words, online TMS applied to a region causally involved in a task may counteract or even reverse the interference effect normally induced by the stimulation, if this region has previously been inhibited by offline 1 Hz TMS (Silvanto et al., 2008a). In this context, the effects observed in our study suggest that, under normal conditions, the posterior cerebellum may send excitatory signals to the pSTS during emotion discrimination, possibly fine-tuning its activity. Consequently, when cerebellar activity is suppressed by offline 1 Hz TMS, the pSTS is indirectly inhibited. Thus, the subsequent online TMS applied over the pSTS results in a facilitatory effect due to this prior cerebellar inhibition.

Accordingly, cerebellar inhibition, which in our study tended to reduce performance when followed by vertex/control online stimulation, has been associated with reduced resting-state functional connectivity between the cerebellum and core regions of the Default Mode Network (DMN), which largely overlaps with the mentalizing network (Rastogi et al., 2017). Thus, our findings may contribute to the ongoing debate about the modulatory tone exerted by the posterior cerebellum on the cerebrum. In fact, while it is widely accepted that the cerebellum exerts an inhibitory tone on the primary motor cortex (M1), a phenomenon known as cerebellar brain inhibition (CBI) (Fernandez et al., 2018), the potential excitatory/inhibitory nature of the posterior cerebellar connectivity with the cerebrum is more controversial. In a previous study, we showed that inhibition of the paravermal posterior cerebellum via TMS reduced motor-evoked potentials during fearful faces viewing (Ferrari et al., 2021). Thus, the direction or tone of the cerebellar modulatory effect (whether inhibitory or excitatory) over the cerebrum may depend on the specific network involved. This perspective is consistent with a previous PET study showing that cerebellar inhibition via 1Hz TMS resulted in both decreased activity in cerebral regions associated with voluntary movements and emotion-related regions, and increased activity in regions associated with cognition and language in the cerebrum (Cho et al., 2012). Overall, the nature of the modulation exerted by the posterior cerebellum on various cortical and subcortical networks warrants further investigation, possibly using double-coils/paired-pulse/cortico-cortical paired associative stimulation methods (Romei et al., 2016).

Finally, it is worth noting that there may be a gradient in posterior cerebellar organization whereby different sectors of the medial/paravermal cerebellum play a role in emotion recognition, whereas lateral sectors are mostly engaged when higher-level inferential processing and mentalizing tasks are tested (Ferrari et al., 2023). This suggests a potentially different time course for the involvement of different posterior cerebellar sectors depending on the specific demands of the task. In our study, we report for the first time that basic social affective processing recruits the cerebellum from approximately 100 ms from stimulus onset in paravermal sectors. We predict that lateral cerebellar regions may be recruited at later stages of processing, when tasks involve higher-level inferential processes such as mentalizing. This is a hypothesis that should be investigated in future studies.

In conclusion, our data are the first to provide insights into the chronometry of social processes occurring in the posterior cerebellum and to show that perturbing cerebellar activity directly affects responses in other nodes of the social brain, such as the pSTS. These findings are critical for a comprehensive understanding of the mechanisms that govern the mentalizing network in the human brain. They also highlight the central role of the cerebellum in social cognition, with important implications for clinical and therapeutic settings.

## Declaration of interests

The authors declare no competing interests.

## Notes

### Competing Interest Statement

The authors have declared no competing interest.

## References

Andersen, L. M., Jerbi, K., & Dalal, S. S. (2020). Can EEG and MEG detect signals from the human cerebellum?. NeuroImage, 215, 116817.

Babo-Rebelo, M., Puce, A., Bullock, D., Hugueville, L., Pestilli, F., Adam, C.,…& George, N. (2022). Visual information routes in the posterior dorsal and ventral face network studied with intracranial neurophysiology and white matter tract endpoints. Cerebral Cortex, 32(2), 342–366.

Battelli, L., Alvarez, G. A., Carlson, T., & Pascual-Leone, A. (2009). The role of the parietal lobe in visual extinction studied with transcranial magnetic stimulation. Journal of cognitive neuroscience, 21(10), 1946–1955.

Battelli, L., Grossman, E. D., & Plow, E. B. (2017). Local immediate versus long-range delayed changes in functional connectivity following rTMS on the visual attention network. Brain stimulation, 10(2), 263–269.

Bijsterbosch, J.D., Barker, A.T., Lee, K.H., Woodruff, P.W. (2012). Where does transcranial magnetic stimulation (TMS) stimulate? Modelling of induced field maps for some common cortical and cerebellar targets. Medical & Biological Engineering & Computing, 50(7), 671–81.

Borgomaneri, S., Zanon, M., Di Luzio, P., Cataneo, A., Arcara, G., Romei, V.,…& Avenanti, A. (2023). Increasing associative plasticity in temporo-occipital back-projections improves visual perception of emotions. Nature Communications, 14(1), 5720.

Buckner, R. L., Krienen, F. M., Castellanos, A., Diaz, J. C., & Yeo, B. T. (2011). The organization of the human cerebellum estimated by intrinsic functional connectivity. Journal of neurophysiology, 106(5), 2322–2345.

Campana, G., Cowey, A., & Walsh, V. (2006). Visual area V5/MT remembers “what” but not “where”. Cerebral Cortex, 16(12), 1766–1770.

Carducci, F., Brusco, R. (2012). Accuracy of an individualized MR-based head model for navigated brain stimulation. Psychiatry Research: Neuroimaging, 203(1), 105–8.

Cattaneo, Z., Bona, S., Ciricugno, A., & Silvanto, J. (2022). The chronometry of symmetry detection in the lateral occipital (LO) cortex. Neuropsychologia, 167, 108160.

Cattaneo, Z., Renzi, C., Casali, S., et al. (2014). Cerebellar vermis plays a causal role in visual motion discrimination. Cortex, 58, 272–80.

Cho, S.S., Yoon, E.J., Bang, S.A., et al. (2012). Metabolic changes of cerebrum by repetitive transcranial magnetic stimulation over lateral cerebellum: a study with FDG PET. The Cerebellum, 11(3), 739–48.

Clausi, S., Lupo, M., Funghi, G., Mammone, A., & Leggio, M. (2022). Modulating mental state recognition by anodal tDCS over the cerebellum. Scientific Reports, 12(1), 22616.

de Graaf, T. A., Koivisto, M., Jacobs, C., & Sack, A. T. (2014). The chronometry of visual perception: review of occipital TMS masking studies. Neuroscience & Biobehavioral Reviews, 45, 295–304.

Demirtas-Tatlidede, A., Freitas, C., Pascual-Leone, A., Schmahmann, J.D. (2011). Modulatory effects of theta burst stimulation on cerebellar nonsomatic functions. Cerebellum, 10(3), 495–503.

Engell, A. D., & Haxby, J. V. (2007). Facial expression and gaze-direction in human superior temporal sulcus. Neuropsychologia, 45(14), 3234–3241.

Ethofer, T., Bretscher, J., Wiethoff, S., Bisch, J., Schlipf, S., Wildgruber, D., & Kreifelts, B. (2013). Functional responses and structural connections of cortical areas for processing faces and voices in the superior temporal sulcus. Neuroimage, 76, 45–56.

Fernandez, L., Major, B.P., Teo, W.P., Byrne, L.K., Enticott, P.G. (2018). The impact of stimulation intensity and coil type on reliability and tolerability of cerebellar brain inhibition (CBI) via dual-coil TMS. The Cerebellum, 17(5), 540–9.

Ferrari, C., Ciricugno, A., Arioli, M., & Cattaneo, Z. (2023). Functional segregation of the human cerebellum in social cognitive tasks revealed by TMS. Journal of Neuroscience, 43(20), 3708–3717.

Ferrari, C., Ciricugno, A., Battelli, L., Grossman, E. D., & Cattaneo, Z. (2022b). Distinct cerebellar regions for body motion discrimination. Social cognitive and affective neuroscience, 17(1), 72–80.

Ferrari, C., Ciricugno, A., Urgesi, C., & Cattaneo, Z. (2022a). Cerebellar contribution to emotional body language perception: a TMS study. Social cognitive and affective neuroscience, 17(1), 81–90.

Ferrari, C., Fiori, F., Suchan, B., Plow, E. B., & Cattaneo, Z. (2021). TMS over the posterior cerebellum modulates motor cortical excitability in response to facial emotional expressions. European Journal of Neuroscience, 53(4), 1029–1039.

Ferrari, C., Oldrati, V., Gallucci, M., Vecchi, T., & Cattaneo, Z. (2018a). The role of the cerebellum in explicit and incidental processing of facial emotional expressions: a study with transcranial magnetic stimulation. NeuroImage, 169, 256–264.

Ferrari, C., Schiavi, S., & Cattaneo, Z. (2018b). TMS over the superior temporal sulcus affects expressivity evaluation of portraits. Cognitive, Affective, & Behavioral Neuroscience, 18, 1188–1197.

Ferrucci, R., Giannicola, G., Rosa, M., Fumagalli, M., Boggio, P. S., Hallett, M.,…& Priori, A. (2012). Cerebellum and processing of negative facial emotions: cerebellar transcranial DC stimulation specifically enhances the emotional recognition of facial anger and sadness. Cognition & emotion, 26(5), 786–799.

Fusar-Poli, P., Placentino, A., Carletti, F., Landi, P., Allen, P., Surguladze, S.,…& Politi, P. (2009). Functional atlas of emotional faces processing: a voxel-based meta-analysis of 105 functional magnetic resonance imaging studies. Journal of psychiatry and neuroscience, 34(6), 418–432.

Gamond, L., Ferrari, C., La Rocca, S., & Cattaneo, Z. (2017). Dorsomedial prefrontal cortex and cerebellar contribution to inlgroup attitudes: a transcranial magnetic stimulation study. European Journal of Neuroscience, 45(7), 932–939.

Guell, X. (2022). Functional gradients of the cerebellum: a review of practical applications. The Cerebellum, 21(6), 1061–1072.

Guell, X., Schmahmann, J. D., Gabrieli, J. D., & Ghosh, S. S. (2018). Functional gradients of the cerebellum. Elife, 7, e36652.

Hanajima, R., Wang, R., Nakatani-Enomoto, S., et al. (2007). Comparison of different methods for estimating motor threshold with transcranial magnetic stimulation. Clinical Neurophysiology, 118, 2120– 2.

Heinen, K., Jolij, J., & Lamme, V. A. (2005). Figure–ground segregation requires two distinct periods of activity in V1: A transcranial magnetic stimulation study. Neuroreport, 16(13), 1483–1487.

Heleven, E., Van Dun, K., De Witte, S., Baeken, C., & Van Overwalle, F. (2021). The role of the cerebellum in social and non-social action sequences: a preliminary LF-rTMS study. Frontiers in Human Neuroscience, 15, 593821.

Ito, M. (2008). Control of mental activities by internal models in the cerebellum. Nature Reviews Neuroscience, 9(4), 304–313.

Kanai, R., Lloyd, H., Bueti, D., & Walsh, V. (2011). Modality-independent role of the primary auditory cortex in time estimation. Experimental Brain Research, 209, 465–471.

Leggio, M., & Molinari, M. (2015). Cerebellar sequencing: a trick for predicting the future. The Cerebellum, 14, 35–38.

Metoki, A., Wang, Y., & Olson, I. R. (2022). The social cerebellum: a large-scale investigation of functional and structural specificity and connectivity. Cerebral Cortex, 32(5), 987–1003.

Oldrati, V., Ferrari, E., Butti, N., Cattaneo, Z., Borgatti, R., Urgesi, C., & Finisguerra, A. (2021). How social is the cerebellum? Exploring the effects of cerebellar transcranial direct current stimulation on the prediction of social and physical events. Brain Structure and Function, 226(3), 671–684.

Pitcher, D. (2014). Facial expression recognition takes longer in the posterior superior temporal sulcus than in the occipital face area. Journal of Neuroscience, 34(27), 9173–9177.

Pitcher, D., & Ungerleider, L. G. (2021). Evidence for a third visual pathway specialized for social perception. Trends in Cognitive Sciences, 25(2), 100–110.

Pitcher, D., Walsh, V., Yovel, G., & Duchaine, B. (2007). TMS evidence for the involvement of the right occipital face area in early face processing. Current Biology, 17(18), 1568–1573.

Rastogi, A., Cash, R., Dunlop, K., Vesia, M., Kucyi, A., Ghahremani, A.,…& Chen, R. (2017). Modulation of cognitive cerebello-cerebral functional connectivity by lateral cerebellar continuous theta burst stimulation. Neuroimage, 158, 48–57.

Renzi, C., Vecchi, T., D’Angelo, E., Silvanto, J., & Cattaneo, Z. (2014). Phosphene induction by cerebellar transcranial magnetic stimulation. Clinical Neurophysiology: Official Journal of the International Federation of Clinical Neurophysiology, 125(10), 2132–2133.

Romei, V., Chiappini, E., Hibbard, P. B., & Avenanti, A. (2016). Empowering reentrant projections from V5 to V1 boosts sensitivity to motion. Current Biology, 26(16), 2155–2160.

Rossi, S., Hallett, M., Rossini, P.M., Pascual-Leone, A. (2011). Screening questionnaire before TMS: an update. Clinical Neurophysiology, 122, 1686–6.

Rossini, P.M., Barker, A.T., Berardelli, A., et al. (1994). Non-invasive electrical and magnetic stimulation of the brain, spinal cord and roots: basic principles and procedures for routine clinical application. Report of an IFCN committee. Electroencephalography and Clinical Neurophysiology, 91(2), 79–92.

Santavirta, S., Karjalainen, T., Nazari-Farsani, S., Hudson, M., Putkinen, V., Seppälä, K.,…& Nummenmaa, L. (2023). Functional organization of social perception networks in the human brain. NeuroImage, 272, 120025.

Schmahmann, J. D., & Pandya, D. N. (1991). Projections to the basis pontis from the superior temporal sulcus and superior temporal region in the rhesus monkey. Journal of Comparative Neurology, 308(2), 224–248.

Schraa-Tam, C. K., Rietdijk, W. J., Verbeke, W. J., Dietvorst, R. C., van den Berg, W. E., Bagozzi, R. P., & De Zeeuw, C. I. (2012). fMRI activities in the emotional cerebellum: a preference for negative stimuli and goal-directed behavior. The Cerebellum, 11, 233–245.

Seitz, R. J., Schäfer, R., Scherfeld, D., Friederichs, S., Popp, K., Wittsack, H. J.,…& Franz, M. (2008). Valuating other people’s emotional face expression: a combined functional magnetic resonance imaging and electroencephalography study. Neuroscience, 152(3), 713–722.

Silvanto, J., Bona, S., & Cattaneo, Z. (2017). Initial activation state, stimulation intensity and timing of stimulation interact in producing behavioral effects of TMS. Neuroscience, 363, 134–141.

Silvanto, J., Bona, S., Marelli, M., & Cattaneo, Z. (2018). On the mechanisms of transcranial magnetic stimulation (TMS): how brain state and baseline performance level determine behavioral effects of TMS. Frontiers in psychology, 9, 741.

Silvanto, J., Cattaneo, Z., Battelli, L., & Pascual-Leone, A. (2008a). Baseline cortical excitability determines whether TMS disrupts or facilitates behavior. Journal of neurophysiology, 99(5), 2725–2730.

Silvanto, J., Muggleton, N., & Walsh, V. (2008b). State-dependency in brain stimulation studies of perception and cognition. Trends in cognitive sciences, 12(12), 447–454.

Sliwinska, M. W., & Pitcher, D. (2018). TMS demonstrates that both right and left superior temporal sulci are important for facial expression recognition. NeuroImage, 183, 394–400.

Sliwinska, M. W., Elson, R., & Pitcher, D. (2020). Dual-site TMS demonstrates causal functional connectivity between the left and right posterior temporal sulci during facial expression recognition. Brain Stimulation, 13(4), 1008–1013.

Sokolov, A. A., Erb, M., Gharabaghi, A., Grodd, W., Tatagiba, M. S., & Pavlova, M. A. (2012). Biological motion processing: the left cerebellum communicates with the right superior temporal sulcus. Neuroimage, 59(3), 2824–2830.

Sokolov, A. A., Erb, M., Grodd, W., & Pavlova, M. A. (2014). Structural loop between the cerebellum and the superior temporal sulcus: evidence from diffusion tensor imaging. Cerebral cortex, 24(3), 626–632.

Sokolov, A. A., Miall, R. C., & Ivry, R. B. (2017). The cerebellum: adaptive prediction for movement and cognition. Trends in cognitive sciences, 21(5), 313–332.

Streit, M., Dammers, J., Simsek-Kraues, S., Brinkmeyer, J., Wölwer, W., & Ioannides, A. (2003). Time course of regional brain activations during facial emotion recognition in humans. Neuroscience letters, 342(1-2), 101–104.

Styliadis, C., Ioannides, A. A., Bamidis, P. D., & Papadelis, C. (2015). Distinct cerebellar lobules process arousal, valence and their interaction in parallel following a temporal hierarchy. Neuroimage, 110, 149–161.

Talairach, J., Tournoux, P. (1998). Co-planar Stereotaxic Atlas of the Human Brain, New York: Thieme Medical.

Thielscher, A., Antunes, A., & Saturnino, G. B. (2015, August). Field modeling for transcranial magnetic stimulation: a useful tool to understand the physiological effects of TMS?. In 2015 37th annual international conference of the IEEE engineering in medicine and biology society (EMBC) (pp. 222-225). IEEE.

Thilo, K. V., Santoro, L., Walsh, V., & Blakemore, C. (2004). The site of saccadic suppression. Nature neuroscience, 7(1), 13–14.

Tottenham, N., Tanaka, J. W., Leon, A. C., McCarry, T., Nurse, M., Hare, T. A.,…& Nelson, C. (2009). The NimStim set of facial expressions: Judgments from untrained research participants. Psychiatry research, 168(3), 242–249.

Tyrrell, R. A., & Owens, D. A. (1988). A rapid technique to assess the resting states of the eyes and other threshold phenomena: the modified binary search (MOBS). Behavior Research Methods, Instruments, & Computers, 20(2), 137–141.

van Dun, K., Bodranghien, F., Manto, M., Mariën, P. (2017). Targeting the cerebellum by noninvasive neurostimulation: a review. Cerebellum, 16(3), 695–741.

Van Overwalle, F., & Mariën, P. (2016). Functional connectivity between the cerebrum and cerebellum in social cognition: a multi-study analysis. Neuroimage, 124, 248–255.

Van Overwalle, F., De Coninck, S., Heleven, E., Perrotta, G., Taib, N. O. B., Manto, M., & Mariën, P. (2019a). The role of the cerebellum in reconstructing social action sequences: a pilot study. Social cognitive and affective neuroscience, 14(5), 549–558.

Van Overwalle, F., Manto, M., Cattaneo, Z., Clausi, S., Ferrari, C., Gabrieli, J. D.,…& Leggio, M. (2020). Consensus paper: cerebellum and social cognition. The Cerebellum, 19, 833–868.

Van Overwalle, F., Van de Steen, F., & Mariën, P. (2019b). Dynamic causal modeling of the effective connectivity between the cerebrum and cerebellum in social mentalizing across five studies. Cognitive, Affective, & Behavioral Neuroscience, 19, 211–223.

Walbrin, J., Downing, P., & Koldewyn, K. (2018). Neural responses to visually observed social interactions. Neuropsychologia, 112, 31–39.

Walsh, V., & Pascual-Leone, A. (2003). Transcranial magnetic stimulation: a neurochronometrics of mind. MIT press.

Weise, K., Numssen, O., Thielscher, A., Hartwigsen, G., & Knösche, T. R. (2020). A novel approach to localize cortical TMS effects. Neuroimage, 209, 116486.

Zhu, Q., Nelissen, K., Van den Stock, J., De Winter, F. L., Pauwels, K., de Gelder, B.,…& Vandenbulcke, M. (2013). Dissimilar processing of emotional facial expressions in human and monkey temporal cortex. Neuroimage, 66, 402–411.

